# A novel RyR2-selective stabilizer prevents stress-induced ventricular arrhythmias without impairing cardiac function

**DOI:** 10.1101/2024.11.26.625386

**Authors:** Nagomi Kurebayashi, Masami Kodama, Takashi Murayama, Masato Konishi, Masami Sugihara, Hana Inoue, Ryosuke Ishida, Koichiro Ishii, Shuichi Mori, Yukari Endo, Xi Zeng, Yukiko U. Inoue, Takayoshi Inoue, Satoru Noguchi, Hajime Nishio, Utako Yokoyama, Junko Kurokawa, Hiroyuki Kagechika, Takashi Sakurai

## Abstract

**Background and Purpose:** Aberrant activation of the type 2 ryanodine receptor (RyR2) causes lethal arrhythmias, such as catecholaminergic polymorphic ventricular tachycardia (CPVT). Developing drugs that suppress RyR2 hyperactivation may be key to novel arrhythmia treatments. This study evaluated the antiarrhythmic potential of Ryanozole, a recently developed novel RyR2 modulator with high affinity and selectivity, using CPVT mouse models harboring mutant RyR2s.

**Experimental approach:** *In vitro* effects of Ryanozole were evaluated by ER Ca^2+^-based assay and [^3^H]Ryanodine binding assay using RyR2-expressing HEK293 cells. Two lines of mice with different arrhythmia severity, RyR2-R420W and -K4750Q, were employed for *in vivo* assessments. Intracellular Ca^2+^ signals were analyzed in isolated cardiomyocytes using Cal520. Antiarrhythmic effects were evaluated by ECG under catecholaminergic challenge in anesthetized mice and during spontaneous arrhythmias in conscious mice. ECG and echocardiographic parameters were evaluated before and after drug administration.

**Key results:** Ryanozole inhibited both wild type and mutant RyR2s with similar IC_50_ of 15−40 nM. The inhibition was more potent at lower cytosolic Ca^2+^ concentrations. It suppressed Ca^2+^ waves and Ca^2+^ sparks without affecting action potential-evoked Ca^2+^ transients. Ryanozole effectively prevented adrenaline-induced arrhythmias and rapidly terminated spontaneous arrhythmias during daily activity. Importantly, Ryanozole did not impair cardiac conduction or contractility, unlike conventional antiarrhythmic drugs.

**Conclusions and implications:** Ryanozole strongly suppresses RyR2 under diastolic Ca^2+^ conditions, thereby preventing the arrhythmogenic trigger of aberrant Ca^2+^ release. This mechanism likely provides potent antiarrhythmic effects while preserving cardiac function. Ryanozole is a promising therapeutic candidate treating RyR2-mediated arrhythmias, such as CPVT.

## Introduction

Type 2 ryanodine receptor (RyR2) is a Ca^2+^ release channel in the sarcoplasmic reticulum (SR) and plays a pivotal role in cardiac muscle excitation–contraction coupling. Genetic variants in the *RYR2* gene, both gain-of-function and loss-of-function ones, are associated with arrhythmogenic heart diseases, including catecholaminergic polymorphic ventricular tachycardia (CPVT) and calcium release deficiency syndrome (CRDS) (Fujii et al., 2017; Hirose et al., 2022; Medeiros-Domingo et al., 2009; Priori & Chen, 2011; Sun et al., 2021; Tester, Spoon, Valdivia, Makielski, & Ackerman, 2004), and more than 300 arrhythmogenic variants have been reported so far. Of these, CPVT is of the gain-of-function type and the most commonly reported diseases caused by *RYR2* variants. The severity of arrhythmias in these variant carriers varies widely: some develop ventricular arrhythmias for the first time in adulthood in response to severe stress, while others suffer frequent arrhythmias from infancy in response to mild exercise or emotional stress (Kawamura et al., 2013). The main cellular mechanism underlying CPVT is as follows: when gain-of-function mutant RyR2 is further activated during strong sympathetic stimulation, spontaneous Ca^2+^ release such as Ca^2+^ waves and Ca^2+^ sparks occur without Ca^2+^ entry from L-type Ca^2+^ channel (LTCC). This in turn activates inward sodium-calcium exchanger (NCX) currents to cause delayed afterdepolarization and triggered activity, leading to arrhythmias (Keefe, Moore, Ho, & Wehrens, 2023; Lakatta, 1992). The severity of the arrhythmia is thought to depend mainly on the degree of enhancement of RyR2 activity caused by each mutation, which determines the feasibility of spontaneous Ca^2+^ release (Kurebayashi et al., 2022).

The major conventional antiarrhythmic drugs for CPVT include β-blockers, the class 1C Na^+^ channel blocker flecainide, and the Ca^2+^ channel blocker verapamil (Bergeman et al., 2023; Priori et al., 2013; van der Werf et al., 2012). Flecainide, which is also reported to normalize RyR2 activity, is effective in CPVT patients (van der Werf et al., 2012; Watanabe et al., 2009), but may decrease cardiac conduction velocity by slowing upstroke of action potentials in atrial and ventricular cells. β-blockers and Ca^2+^ channel blockers can suppress atrioventricular conduction and weaken cardiac contractility. Implantable cardioverter defibrillators (ICDs) are recommended in severe cases, but patients with ICDs are at risk of receiving inappropriate shocks, which may worsen the arrhythmias (Itoh et al., 2021; Lamba et al., 2024). Thus, developing drugs with sufficient efficacy and fewer adverse effects to treat CPVT caused by various *RYR2* variants would be beneficial. Furthermore, it is desirable for the drugs to immediately stop already occurring arrhythmias in addition to prevent future arrhythmias.

Based on the pathogenic mechanisms of CPVT caused by hyperactivated RyR2, attempts have been made to find drugs that suppress RyR2 activity. Several drugs and compounds have been proposed to suppress RyR2, which include flecainide, dantrolene, carvedilol, EL20, and benzothiazepine derivatives S107 and K201, and unnatural verticilide enantiomers (Batiste et al., 2019; Do & Knollmann, 2025; Lehnart et al., 2008; Szentandrassy et al., 2022; Watanabe et al., 2009; Zhou et al., 2011). Recently, we established an effective platform for screening RyR2 modulators using HEK293 cells expressing RyR2 and ER Ca^2+^ sensor protein R-CEPIA1er (Murayama & Kurebayashi, 2019; Murayama et al., 2018; Takenaka et al., 2023). We identified several compounds that selectively modulate RyR2 activity without affecting RyR1 or RyR3 (Ishida et al., 2023; Takenaka et al., 2023). In addition, we developed a series of tetrazole compounds (Ryanozoles) that selectively stabilize RyR2 in the closed state (Ishida et al., 2023). Among these Ryanozoles, TMDJ-035, which we will refer to here as Ryanozole, had the highest affinity for RyR2 with IC_50_ of 15 nM (Ishida et al., 2023). However, the *in vivo* effects of ryanozole, i.e., how effectively it can suppress arrhythmias, have not yet been proven. In addition, it has not been investigated whether ryanozole’s RyR2 inhibition has any adverse effects on cardiac function.

In this study, we investigated the *in vitro* and *in vivo* effects of Ryanozole using two lines of mice with CPVT-associated RyR2 mutations showing different disease severity. We found that Ryanozole effectively suppresses Ca^2+^ sparks and Ca^2+^ waves without affecting action potential-evoked Ca^2+^ transients in the isolated myocytes. Importantly Ryanozole prevented induced and spontaneous arrhythmias in these mice without affecting parameters of ECG and echocardiogram. These preferred effects were caused by the unique property of Ryanozole that suppresses RyR2 more strongly at low cytoplasmic Ca^2+^ concentrations ([Ca^2+^]_cyt_) than that at high [Ca^2+^]_cyt_, Our results provide the strong evidence that Ryanozole should be a promising therapeutic candidate for the treatment of CPVT without severe side effects.

## Methods

### ER Ca^2+^-based assay using HEK293 cells

HEK293 cells stably expressing ER Ca^2+^ sensor protein R-CEPIA1er (Suzuki et al., 2014) together with inducibly expressing RyR2 wild type (WT) were generated as described previously (Murayama et al., 2018; Takenaka et al., 2023). Cells were cultured in Dulbecco’s Modified Eagle’s Medium supplemented with 10% fetal calf serum, 2 mM L-glutamine, 15 µg/ml blasticidin, 100 µg/ml hygromycin, and 400 µg/ml G418. To test the effects of compounds on mutant RyR2s, RyR2-R420W, -K4750Q, R2474S, R4497C, phospho-null triple mutant S2807A/S2813A/S2030A (RyR2-S3A), and phospho-mimetic triple mutant (RyR2-S3D) cells (Kurebayashi et al., 2022; Takenaka et al., 2023; Uehara et al., 2017), these cells were infected with baculovirus vector to express R-CEPIA1er.

Time-lapse [Ca^2+^]_ER_ measurements were performed using the FlexStation3 fluorometer (Molecular Devices, San Jose, CA, USA), as described previously (Takenaka et al., 2023). Briefly, HEK293 cells expressing R-CEPIA1er and RyR2 WT on 96-well, flat, clear-bottomed black microplates (#3603; Corning, New York, NY, USA) were used in this study. Before the measurements, the culture medium in individual wells was replaced with 90 µl of HEPES-Krebs solution (140 mM NaCl, 5 mM KCl, 2 mM CaCl_2_, 1 mM MgCl_2_, 11 mM glucose, and 10 mM HEPES, pH 7.4). R-CEPIA1er was excited at 560 nm and fluorescence emitted at 610 nm was captured every 10 s for 260 s. At 60 s after the start of measurement, 60 µl of compound solution, containing 25 µM test compound in HEPES-Krebs solution, was applied to the wells of the reading plate at a final concentration of 0.001–10 µM. The change in fluorescence induced by the compounds was expressed as F/F_0_, in which mean fluorescence intensity of the last 60 s (F) was normalized to that of the initial 60 s (F_0_). Measurements were performed at 37°C.

### [^3^H]Ryanodine binding assay

[^3^H]Ryanodine binding was carried out as described previously (Kurebayashi, et al, 2022; Takenaka et al, 2023). Microsomes isolated from the HEK293 cells expressing RyR2 or mouse hearts were incubated for 1 h at 25°C with 5 nM [^3^H]ryanodine in a medium containing 0.17 M NaCl, 20 mM 3-(N-morpholino)-2-hydroxypropanesulfonic acid (MOPSO) at pH 7.0, 2 mM dithiothreitol, 1 mM AMP, 1 mM MgCl_2_, and various concentrations of free Ca^2+^ buffered with 10 mM ethylene glycol-bis(2-aminoethylether)-N,N,N,N-tetraacetic acid (EGTA). Free Ca^2+^ concentrations were calculated using the WEBMAXC STANDARD (https://somapp.ucdmc.ucdavis.edu/pharmacology/bers/maxchelator/webmaxc/webmaxcS.htm). The protein-bound [^3^H]ryanodine was separated by filtering through polyethyleneimine-treated GF/B filters using Micro 96 Cell Harvester (Skatron Instruments). Nonspecific binding was determined in the presence of 20 µM unlabeled ryanodine. [^3^H]Ryanodine-binding data (*B*) were normalized to the maximum number of functional channels (*B*_max_), which was separately determined by Scatchard plot analysis using various concentrations (3–20 nM) of [^3^H]ryanodine in a high-salt medium.

### Animals

All animal experiments were approved by the Institutional Animal Care and Use Committee of Juntendo University and NCNP, and were performed in accordance with the guidelines of the Laboratory Animal Experimentation of the Juntendo University and NCNP, both established based on the NIH Guide for the Care and Use of Laboratory Animals. Mice were bred in dedicated breeding facilities, provided with food and water ad libitum, and kept in a controlled environment with a 12/12 h light/dark cycle, temperature of 23–25°C, and relative humidity of 50%–60% in specific pathogen-free conditions at Juntendo University.

### Generation of RYR2-p.K4750Q mice using the CRISPR-Cas9 gene editing system

The mouse genomic sequence within 50 bp both upstream and downstream of each human mutation site was searched by CRISPOR (http://crispor.gi.ucsc.edu/) to select PAM and its consequent guide sequence with high specificity. TGAGGACAGGATAGTTCTCA was selected for crRNA synthesis. A single-stranded donor oligonucleotide carrying K4750Q mutation, FspI site, and modification at PAM to prevent re-editing, TGACCATGTCTGTTCTTGGACATTATAACAACTTTTTTTTTGCTGCTCACCTCCTTGACATCGCGATGGGATTCCAGACATTGCGCACTATCCTGTCCTCAGTTACCCATAATGGCAAACAGGTAAAAGGTCCCATTTCCCTTAGAGAAACTAAAACTGA, was chemically synthesized for homologous recombination. An RYR2-genomic fragment containing PAM and the consequent guide sequence were amplified by PCR from WT mouse genomic DNA. RNP complex formed by the mixing of tracrRNA and crRNA with recombinant Cas9 protein was added to the PCR fragment and incubated at 37°C for 1 h. Cleavage of the PCR fragment into two fragments was confirmed by agarose electrophoresis. RNP complex and single-stranded donor oligonucleotide were introduced to mouse one-cell-stage zygotes by electroporation. The electroporated zygotes were transferred to the oviducts of pseudopregnant females. Genomic DNA was prepared from newborns’ tail and knock-in mice were screened by PCR-RFLP using a pair of primers (forward: CAGCTTAGACCGAGGAGGTG, reverse: AGCTGGCTGGTTGATTTGTT) and Fsp I digestion. The obtained knock-in founders were crossed with WT C57BL/6J mice to obtain the F_1_ generation in line with the standard protocol. Basically, offspring obtained by mating heterozygous and WT C57BL/6J mice were used in the experiments. Only when the properties of homozygous mice were examined, offspring were obtained by mating heterozygous mice.

### Preparation of single cardiomyocytes and Ca^2+^ imaging

WT and mutant mice (R420W and K4750Q, 8–12 weeks old) were deeply anesthetized with pentobarbital sodium (200 mg/kg i.p.) after acute anesthesia with 5 % isoflurane, and ventricular cardiomyocytes were isolated as previously reported (Shioya, 2007; Takenaka et al., 2023). Isolated cardiomyocytes were seeded on a laminin-coated cover slip of the 35 mm glass-bottom dish for Ca^2+^ imaging and loaded with Cal520-AM (AAT Bioquest, Inc., Pleasanton, CA, USA). Cells were field-stimulated at 0.5 Hz or 0.33 Hz in HEPES-buffered Tyrode’s solution (140 mM NaCl, 5 mM KCl, 1.5 mM CaCl_2_, 1 mM MgCl_2_, 0.3 mM NaH_2_PO_4_, 11 mM glucose, 10 mM HEPES, pH7.4) with or without test compounds. Cal520 was excited at 488 nm through a 60× (for line-scan images, Figs. 2C and 5C) or 20× (for mean fluorescence intensities in the ROIs, Figs. 2B, 5B, and 9B) objective lens and light emitted at 525 nm was captured with an EM-CCD camera at 5.3–5.7 ms intervals (Model 8509; Hamamatsu Photonics). Fluorescence signals (F) of Cal520 in individual cells were determined using region of interest (ROI) analysis, from which the cell-free background fluorescence was subtracted. F values were then normalized to the basal levels (F_0_) to yield F/F_0_. For the detection of Ca^2+^ waves and Ca^2+^ sparks, time-based scan images generated by ImageJ software (Schneider, Rasband, & Eliceiri, 2012) were analyzed with SparkMaster 2 (Tomek, Nieves-Cintron, Navedo, Ko, & Bers, 2023). All measurements were carried out at 30°C by perfusing solutions through an inline solution heater/cooler (Warner Instruments, Holliston, MA, USA).

### Patch-clamp recordings

Simultaneous recording of action potentials and Ca^2+^ transients in cardiomyocytes was performed at 32±1°C. The instruments and procedures for the recordings were as described previously (Okabe et al., 2024). Briefly, action potentials were triggered by a 1 ms depolarizing current injection at 1 Hz 30 times under current clamp mode. The intracellular fluorescence signals of Fluo-4 excited with 490 nm were collected at 3.9 ms sampling intervals. Cardiomyocytes were superfused with a solution containing 135 mM NaCl, 4.0 mM KCl, 1.2 mM NaH_2_PO_4_, 1.0 mM MgCl_2_, 10 mM HEPES, 10 mM glucose, and 1.8 mM CaCl_2_, adjusted to pH 7.4 with NaOH. The pipette solution contained 120 mM K-aspartate, 20 mM KCl, 1.0 mM MgCl_2_, 4.0 mM Na_2_ATP, 0.1 mM Na_2_GTP, 10 mM HEPES, and 0.2 mM Fluo-4 pentapotassium salt, adjusted to pH 7.2 with KOH. When studying the effect of Ryanozole, it was included in both the superfusing solution and the pipette solution at a concentration of 100 nM.

### Reagents

Ryanozole was synthesized in accordance with our reported procedure (Ishida et al., 2023). For *in vitro* experiments, Ryanozole was dissolved in dimethyl sulfoxide (DMSO) and diluted 1000 times in Krebs or Tyrode’s solution. Flecainide acetate salt, (+/−)-verapamil hydrochloride, and atenolol were all purchased from Sigma.

### *In vivo* experiments Drug administration

Ryanozole powder was mixed with DMSO and suspended in 0.5% methylcellulose with a final DMSO concentration of 2%. Because Ryanozole is poorly soluble in aqueous solution, the methylcellulose suspension was vortexed for 30 s and then sonicated for 30 s before administration to mice. Flecainide, verapamil, and atenolol were also administered in the same way as Ryanozole.

### Electrocardiographic analysis

Surface electrocardiography (ECG) was recorded from male and female mice anesthetized with isoflurane (3% for induction, 1.5% for maintenance) using a PowerLab system (AD Instruments, Dunedin, New Zealand) on a heated pad. Electrodes were attached to the mouse paws to record the Lead-II ECG. In experiments comparing drug effects, littermates were randomly assigned to different drug or dose groups, and each group contained approximately equal numbers of male and female mice. The procedure for examining the preventive effects of drugs against catecholamine-induced arrhythmias is described in the results section. The ECG recordings during 30 min after Ad/Caf administration were classified into four types, namely, normal rhythm, T-wave alternans, ventricular bigeminy, and ventricular tachycardia, and their proportions were determined. T-wave alternans is defined as a phenomenon in which the RR and PR intervals are regular, but the amplitude or morphology of the T-wave alternates every other beat. To examine the effects of drugs on ECG parameters, ECG records were analyzed with ECG Analysis software (AD Instruments).

Telemetric ECG recordings were obtained in male K4750Q mice at 3–5 months of age using an implantable telemetry system with ECG transmitters (ETA-F10; Data Sciences International, St. Paul, MN, USA), which was implanted subcutaneously in mice at 2–2.5 months of age. Vehicle, Ryanozole, or conventional drugs (flecainide, atenolol, verapamil) was injected in the evening, between 7 and 8 p.m. Individual mice were administered a series of drugs (various doses of Ryanozole or a set of dugs including vehicle, Ryanozole, flecainide, atenolol and verapamil) in a random order within 3 weeks, with at least 48 hours between drug administrations. The proportions of the time during which ventricular arrhythmias (ventricular bigeminy and ventricular tachycardia) occurred were determined.

### Echocardiography

Transthoracic echocardiography was carried out with a Vevo 2100 instrument (Fujifilm Visual Sonics). Mice were anesthetized with isoflurane (3% for induction, 1% for maintenance) and maintained on a heated pad. Long-axis, short-axis, and four-chamber view images were acquired. The following echocardiographic parameters were measured on M-mode images at the tip of the papillary muscles: interventricular septum thickness at end-diastole (IVS), LV posterior wall thickness at end-diastole (PW), LV diameter at end-diastole (LVDD) and end-systole (LVDS), fractional shortening (FS), and ejection fraction (EF). E/A ratios were evaluated on four-chamber views.

### Statistics

Data are presented as mean±SD. Unpaired Student’s t test was used for comparisons between two groups. One-way or two-way analysis of variance (ANOVA), followed by Tukey’s test, was performed to compare three or more groups. Two-tailed tests were used for all analyses. Statistical analysis was performed using Prism v10 (GraphPad Software, Inc., La Jolla, CA, USA). A P-value□<□0.05 was considered significant; *P□<□0.05; **P□<□0.01; ***P□<□0.005; ****P□<□0.001; ns: not significant. Symbols always indicate comparisons between the control (WT or vehicle-treated) and the other tested conditions.

## Results

### Ryanozole stabilizes various mutant RyR2s in the closed state

Figure 1A shows the chemical structure of Ryanozole TMDJ-035 (denoted here as “Ryanozole”), which selectively suppresses RyR2 among the three RyR subtypes (Ishida et al., 2023). Prior to evaluation in mouse models, we further characterized the effect of Ryanozole on RyR2. First, we tested its effects on the wild type (WT) and various RyR2 mutants using an ER Ca^2+^-based assay (Murayama et al., 2018; Takenaka et al., 2023). In this assay, we measured ER Ca^2+^ signals in HEK293 cells expressing RyR2 and R-CEPIA1er, a fluorescent ER Ca^2+^ sensor protein, and obtained ratios (F/F_0_) of fluorescence intensity before (F_0_) and after addition of the compounds (F) as an indicator of RyR2 inhibition (Fig. S1A, B). Ryanozole dose-dependently increased the F/F_0_ ratio with IC_50_ of 16.5 nM in WT cells (Fig. 1B) and in CPVT-linked mutant cells R420W, K4750Q, R2474S, and R4497C with IC_50_s of 20.2–42.5 nM (Fig. S1C−F). We also examined the effects of Ryanozole on RyR2s carrying a phospho-null triple mutation (S2807A/S2813A/S2030A) (RyR2 S3A) or a phospho-mimetic triple mutation (S2807D/S2813D/S2030D) (RyR2 S3D) at the three major phosphorylation sites (Huke & Bers, 2008; Lanner, Georgiou, Joshi, & Hamilton, 2010; Potenza et al., 2019; Takenaka et al., 2023), as RyR2 has been reported to undergo post-translational modifications such as phosphorylation. The effect of Ryanozole was indistinguishable between the two mutant RyR2s, suggesting its phosphorylation-independent effect (Fig. S1G, H). These results suggest that Ryanozole stabilizes RyR2 in the closed state regardless of the presence of disease-associated mutations or phosphorylation at major phosphorylation sites.

**Fig. 1.**
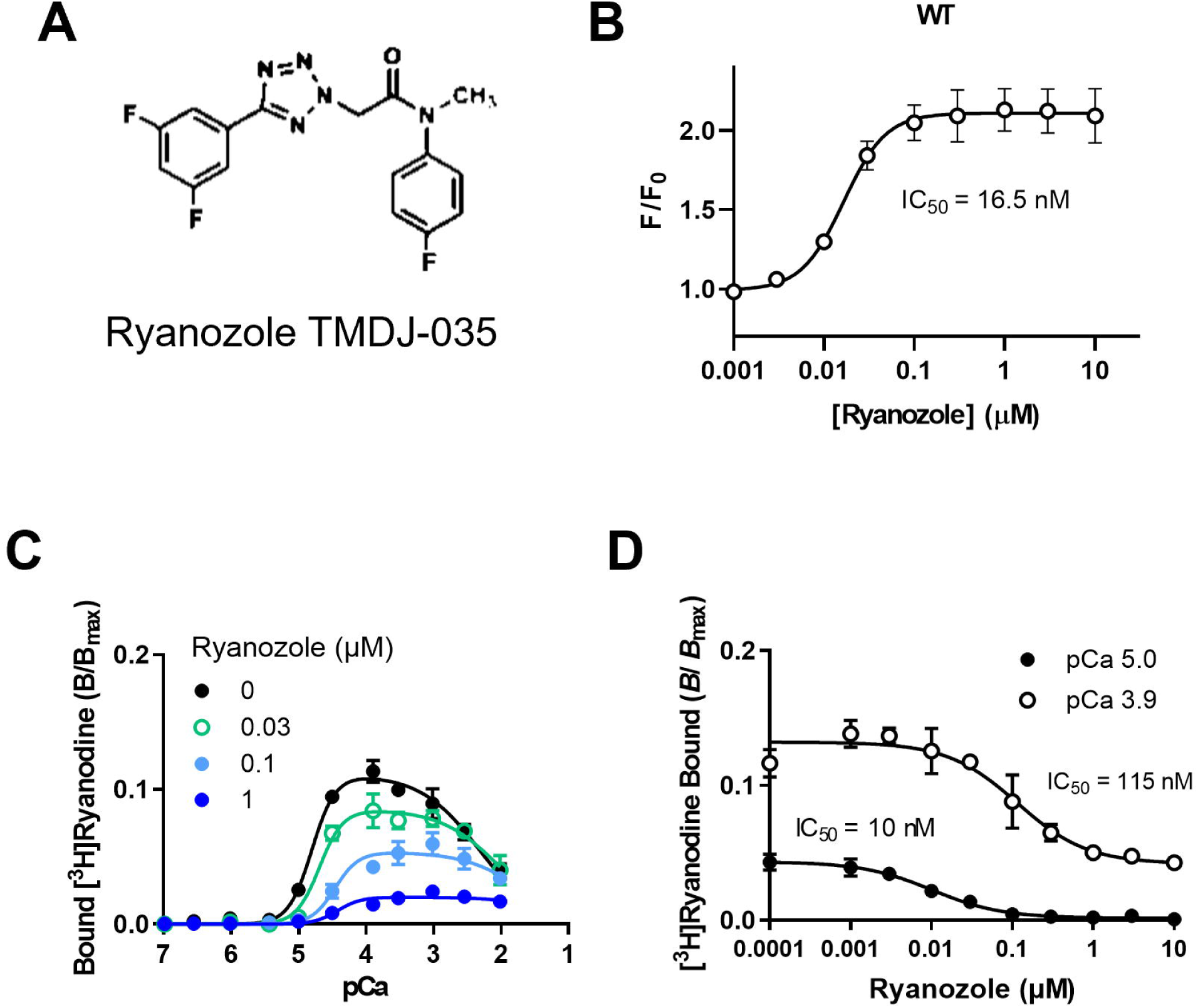
Effects of Ryanozole on RyR2 activity. **A**, Chemical structure of Ryanozole TMDJ-035. **B**, Dose-dependent effects of Ryanozole on ER Ca^2+^ signals in HEK293 cells expressing WT RyR2 determined with FlexStation3. Data are mean±SD (n=5). **C and D**, Effects of Ryanozole on [^3^H]ryanodine binding. Ca^2+^-dependent [^3^H]ryanodine binding in the presence of 0.03, 0.1 and 1 μM Ryanozole (**C**). Dose-dependent effects of Ryanozole at pCa 5.0 (filled circles) and pCa 3.9 (open circles) (**D**). Data are mean±SD (n=4).

### Ryanozole more strongly suppresses RyR2 activity at low [Ca^2+^]_cyt_

The inhibitory effect of Ryanozole on RyR2 was further investigated by [^3^H]ryanodine binding assay, which provides a useful measure of the RyR activity (Fujii et al., 2017; Kurebayashi et al., 2022). Ryanozole reduced the peak of Ca^2+^-dependent [^3^H]ryanodine binding of RyR2 and shifted the Ca^2+^ dependence to the right (Fig. 1C). Dose dependence of Ryanozole at pCa 5.0 indicates that the IC_50_ is 10 nM (Fig. 1D), which is close to that obtained with cell-based assay (Fig. 1B). Interestingly, IC_50_ at pCa 3.9 is 10 times greater (115 nM) than that at pCa 5.0, suggesting that the suppressive effect of Ryanozole was much less marked at higher Ca^2+^ concentrations. Taken together, these results suggest that Ryanozole suppresses the RyR2 activity more strongly at low cytoplasmic Ca^2+^ concentrations ([Ca^2+^]_cyt_) near resting levels than at high [Ca^2+^]_cyt_ during excitation-contraction coupling.

### Two RyR2 mutations for CPVT mouse models

Previous studies indicate that CPVT patients with mild to moderate gain-of-function *RYR2* may develop infrequent ventricular arrhythmias upon intense exercise or emotional stress. In contrast, patients with strong gain-of-function *RYR2* develop frequent arrhythmias from a young age upon moderate daily exercise (Kawamura et al., 2013; Kurebayashi et al., 2022). Antiarrhythmic drugs are expected to serve two main purposes in treating these arrhythmias: preventing future episodes and suppressing ongoing ones. For the latter, the drug must act rapidly. To evaluate these effects, we selected two mouse lines with different arrhythmia severities, RyR2-R420W and -K4750Q, based on *in vitro* measurements using RyR2-expressing HEK293 cells (Kurebayashi et al., 2022). As shown in Fig. S2A, RyR2 activity determined with [^3^H]ryanodine binding assay was moderately increased in R420W, whereas it was more enhanced in K4750Q in microsomes obtained from HEK293 cells expressing homozygous RyR2.

### Ryanozole suppresses abnormal Ca^2+^ release events in cardiomyocytes isolated from RyR2-R420W mice

An RyR2-R420W mouse model was previously created based on cases of sudden unexplained deaths in 13- and 33-year-old victims (Nishio, Iwata, & Suzuki, 2006; Nishio et al., 2014). Arrhythmias have been reported to be induced by the injection of a mixture of adrenaline (2 mg/kg) and caffeine (120 mg/kg), and the frequency of arrhythmias was found to be higher in homozygous than in heterozygous mice (Okudaira, Kuwahara, Hirata, Oku, & Nishio, 2014). This difference in arrhythmia susceptibility can be well explained by differences in RyR2 activity as determined from [^3^H]ryanodine binding in microsomes prepared from mouse ventricles (Fig. S2B, left). We used homozygous R420W mice (hereafter referred to as “R420W mice”) in this study.

We first tested the in vitro effect of Ryanozole on R420W mice using isolated cardiomyocytes. It is well known that abnormal Ca^2+^ release events such as Ca^2+^ waves lead to arrhythmias in CPVT. In R420W cells regularly stimulated at 0.5 Hz, many abnormal Ca^2+^ signals were induced by 100 nM isoproterenol. The addition of Ryanozole at 30 and 100 nM effectively reduced these abnormal signals (Fig. 2A). In a total of 74–132 cells (pooled data from the previous study and the present study), the mean rate of occurrence of abnormal Ca^2+^ signals was 0.11±0.39 events·s^−1^·cell^−1^ with isoproterenol, which was significantly reduced to 0.02±0.13 events·s^−1^·cell^−1^ and to 0.003±0.018 events·s^−1^·cell^−1^ by 30 nM and 100 nM Ryanozole, respectively (Fig. 2B). We also performed “line-scan imaging analysis” using time-based scan images obtained from time-lapse 2D imagings to investigate whether Ryanozole can suppress different types of Ca^2+^ release events, especially Ca^2+^ sparks, which are hardly detected by whole-cell fluorescence intensity (see Methods). In the cells in which frequent Ca^2+^ sparks were induced by isoproterenol (Fig 2C, D) (Video S1), Ryanozole (100 nM) almost completely abolished Ca^2+^ sparks as well as Ca^2+^ waves and triggered activities (Fig. 2C, D) (Video S2).

**Fig. 2.**
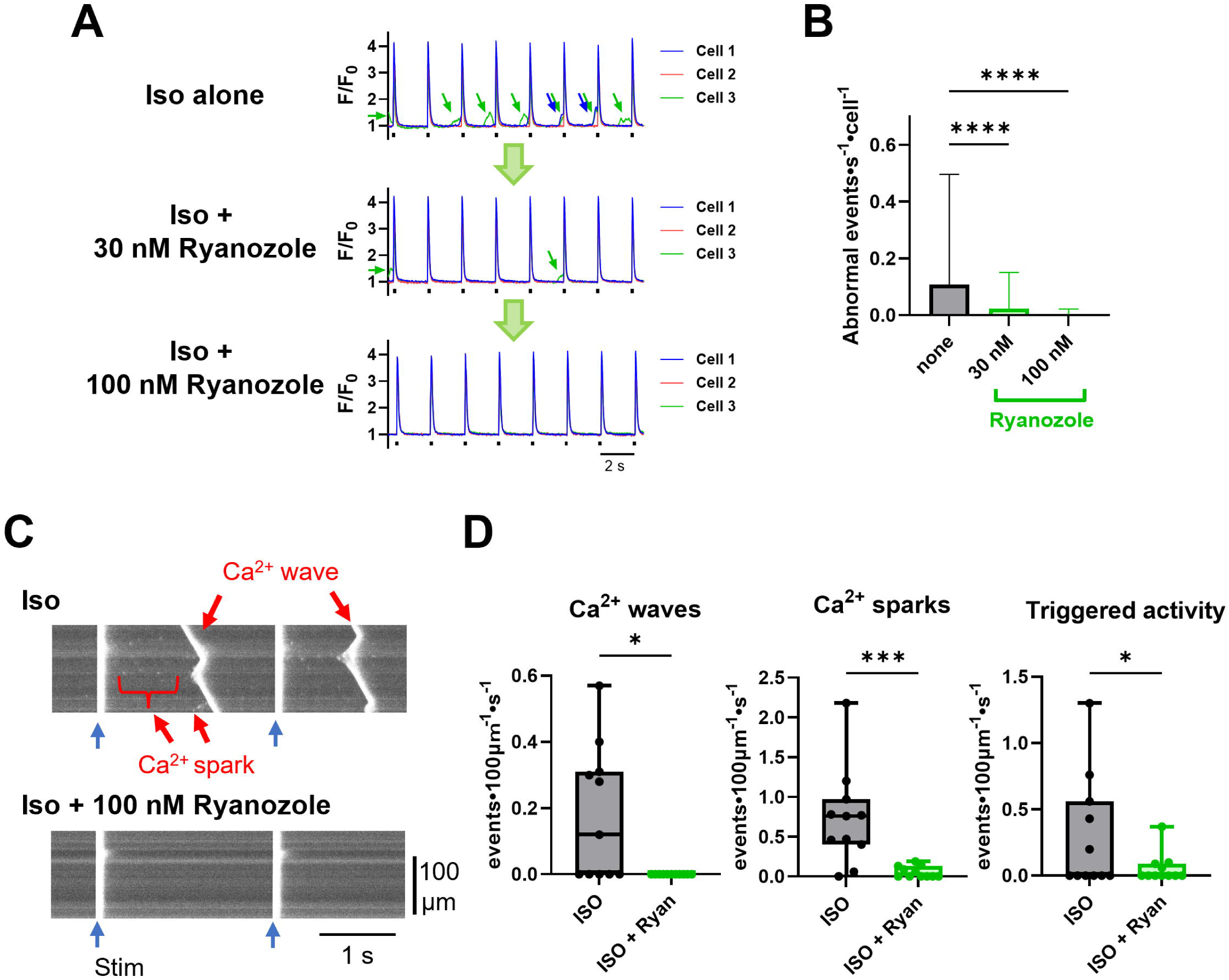
Effects of Ryanozole on spontaneous Ca^2+^ release in R420W cardiomyocytes. **A**, Typical effects of Ryanozole on isoproterenol (Iso)-induced abnormal Ca^2+^ signals (average fluorescence intensities in ROIs in images captured through a 20× objective lens). Cardiomyocytes were field stimulated at 0.5 Hz (indicated with ticks). Arrows indicate Ca^2+^ waves. **B**, Average number of abnormal Ca^2+^ release events before and after the application of Ryanozole. Data are represented as mean±SD (n=74-132, N=8). **** P<0.0001. Statistical significance was analyzed by one-way ANOVA followed by Kruskal−Wallis test. **C**, Typical time-based line scan images obtained with a 60× objective lens before (top) and after (bottom) Ryanozole application. **D**, Effects of 100 nM Ryanozole on frequencies of Ca^2+^ waves, Ca^2+^ sparks and triggered activity (n=11, N=3). Statistical significance was analyzed by Mann−Whitney test. * p < 0.05 and *** P < 0.001 compared to ISO alone.

### Ryanozole prevents induced arrhythmias in R420W mice

We next tested the *in vivo* effect of Ryanozole on the induced arrhythmias using R420W mice. Isoflurane-anesthetized R420W mice aged between 8 and 12 weeks were subjected to an adrenaline and caffeine stress test based on a previous report (Okudaira et al., 2014). In individual animals, a baseline ECG was recorded for 10 min, followed by drug or vehicle injection; 10 min later an adrenaline (2 mg/kg)–caffeine (120 mg/kg) (Ad/Caf) mixture was injected, and the ECG was recorded for another 30 min. Typical ECG records are shown in Fig. 3A. In the vehicle-pretreated mouse, sustained ventricular arrhythmias were frequently observed (Fig. 3A, top). In the mouse pretreated with 30 mg/kg Ryanozole, such ventricular arrhythmias were largely suppressed (Fig. 3A, bottom). We classified ECG waveforms into four types: normal rhythm, T-wave alternans, bigeminy, and ventricular tachycardia (VT) (Fig. 3B), where bigeminy and VT are ventricular arrhythmias. Although T-wave alternans is not an arrhythmia itself, it is considered a pre-arrhythmic marker and has been frequently observed in CPVT patients and mouse models (Kulkarni et al., 2019). The proportion of time for each rhythm/arrhythmia during 30 min after injection was determined for individual mice and the mean values are plotted in Fig. 3C. Severe arrhythmias such as VT were frequently observed in the vehicle-pretreated mice. Ryanozole dose-dependently suppressed the proportion of VT at 3 mg/kg or higher (Fig. 3C). The proportion of normal rhythm was significantly higher at 30 mg/kg Ryanozole.

**Fig. 3.**
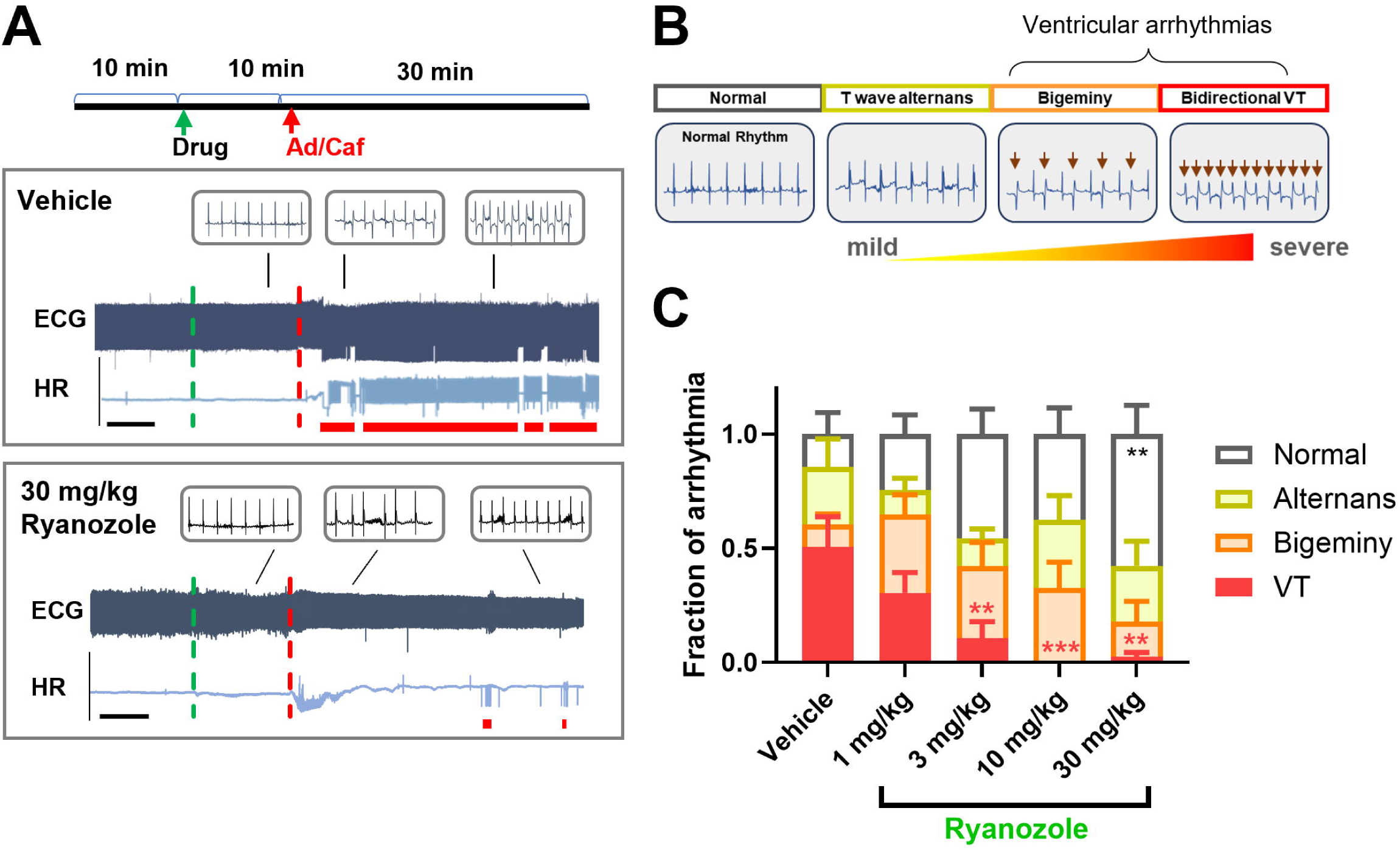
Effect of Ryanozole on catecholamine-induced arrhythmias in R420W mice. **A**, Typical effects of preinjected vehicle (top) and Ryanozole (bottom) on Ad/Caf-induced arrhythmias. Administrations of vehicle/drug and Ad/Caf mixture are indicated by the vertical green and red dotted lines, respectively. Ventricular arrhythmias are indicated with horizontal thick red line. Note that when ventricular arrhythmias occur, the HR traces become abnormal, often appearing as thick bands. Bars indicate 5 min. **B**, Classification of cardiac rhythms. ECG records were classified into four types: normal, T wave alternans, bigeminy, and bidirectional ventricular tachycardia (VT). **C**, Dose-dependent effects of preinjected Ryanozole on proportion of arrhythmias during 30 min after injection of Ad/Caf (N=10). Data are represented as mean±SEM. *P<0.05, **P<0.01, ***P<0.001 compared with Vehicle. Statistical significance was analyzed by two-way ANOVA followed by Dunnett’s multiple comparison test.

### Creation of K4750Q mice

We newly created a knock-in mouse carrying the K4750Q mutation as a severe CPVT model for the evaluation of Ryanozole (Fig. S3A). The heterozygous *RYR2*-K4750Q variant has been reported to cause frequent polymorphic ventricular tachycardia in a young, 6-year-old carrier (Kawamura et al., 2013) and to greatly promote RyR2 activity in an *in vitro* expression system (Kurebayashi et al., 2022; Uehara et al., 2017). Heterozygous RyR2-K4750Q mice were born and grew normally. In contrast, homozygous RyR2-K4750Q mice were born very rarely (3% of the total litters in heterozygous mating) (Fig. S3B) and, even when they were born, they were small and mostly died before the age of 3 months. [^3^H]Ryanodine binding in microsomes from heterozygous K4750Q mouse ventricles was substantially higher than that from homozygous R420W mice (Fig. 4A, see also Fig. S2B), although that from the homozygous K4750Q ventricle was the highest (Fig. S2B, right). The body weight, heart to body weight ratio (HW/BW), and lung to body weight ratio (LW/BW) of heterozygous K4750Q mice were not significantly different from those of WT and R420W (Fig. S3C) and basal ECG parameters (PR interval, QRS interval, and heart rate) were normal (Fig. S3D), consistent with the clinical consensus that typical CPVT is not associated with cardiac hypertrophy or heart failure (Priori et al., 2001; Priori et al., 2013). We used heterozygous K4750Q mice (hereafter referred to as “K4750Q mice”) as another CPVT mouse model to evaluate the antiarrhythmic effects of drugs.

**Fig. 4.**
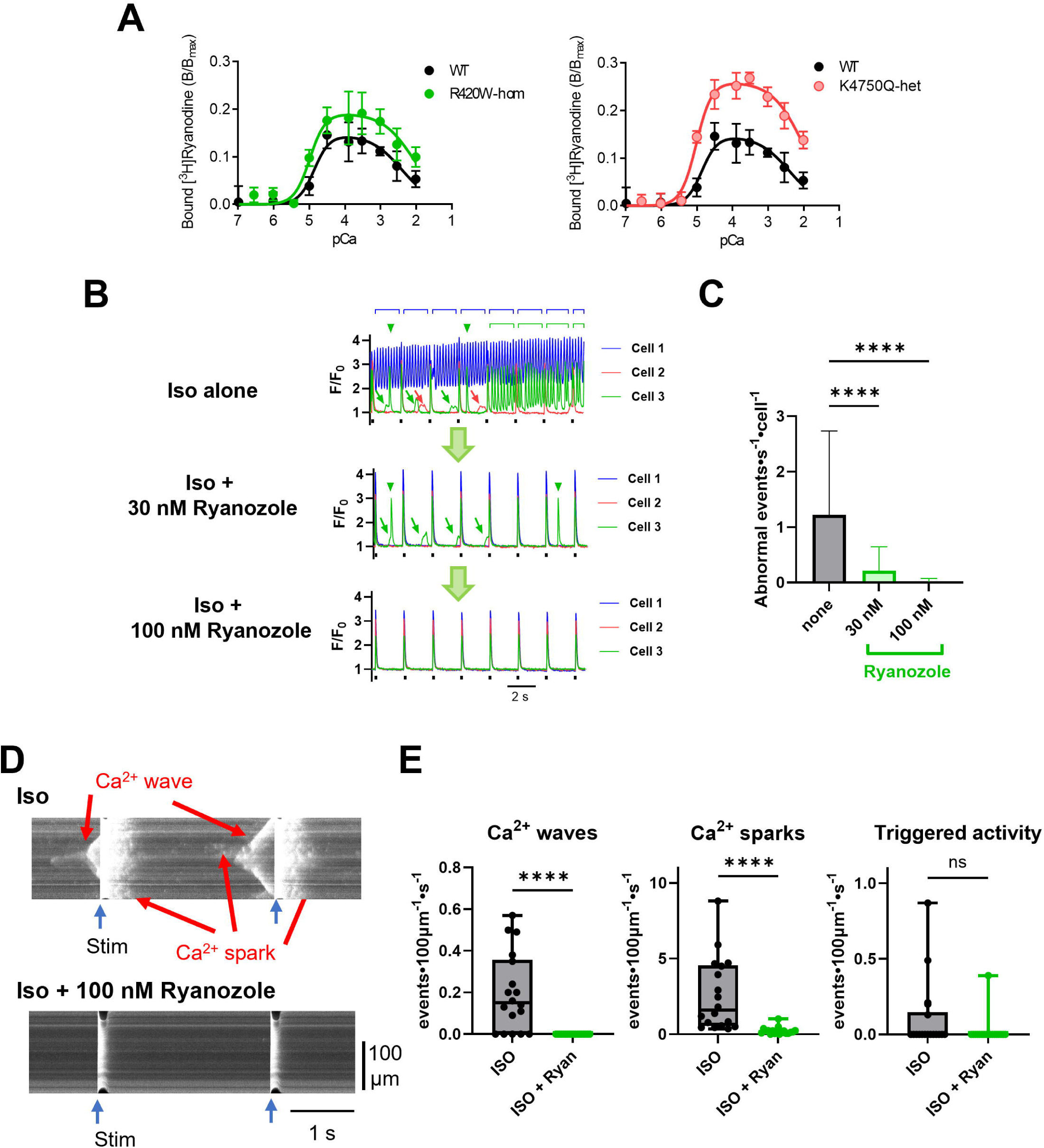
RyR2 activity of myocardium from K4750Q mice and effects of Ryanozole on spontaneous Ca^2+^ release in K4750Q cardiomyocytes. **A**, RyR2 activity determined with [3H]ryanodine binding in ventricular microsomes from heterozygous K4750Q mice (right). [^3^H]Ryanodine binding in ventricular microsomes from homozygous R420W mice is also shown for comparison (left). **B**, Typical effects of Ryanozole on isoproterenol (Iso)-induced abnormal Ca^2+^ signals detected from ROIs. Cardiomyocytes were field stimulated at 0.5 Hz (indicated with ticks). Arrows and bars indicate Ca^2+^ waves. Arrowheads indicate triggered activities. **C**, Average number of abnormal Ca^2+^ release events before and after the application of Ryanozole (n=83, N=4). Data are represented as mean±SD. **** P<0.0001. Statistical significance was analyzed by one-way ANOVA followed by Kruskal−Wallis test. **D**, Typical time-based line scan images before (top) and after (bottom) the Ryanozole application. **E**, Effects of 100 nM Ryanozole on frequencies of Ca^2+^ waves, sparks and triggered activities (n=18, N=2). Statistical significance was analyzed by Mann−Whitney test. * p < 0.05 and *** P < 0.001.

### Ryanozole suppresses abnormal Ca^2+^ release events in cardiomyocytes isolated from K4750Q mice

We determined the *in vitro* effect of Ryanozole on the K4750Q mice using isolated cardiomyocytes. In the presence of 100 nM isoproterenol, more than 90% of RyR2-K4750Q cells were spontaneously active with abnormal Ca^2+^ signals, which was a higher rate than for R420W cells (Fig. 4B). The mean rate of occurrence of abnormal Ca^2+^ signals was 1.23±1.51 events·s^−1^·cell^−1^ with 100 nM isoproterenol (Fig. 4C). Ryanozole effectively suppressed the abnormal Ca^2+^ signals in a dose-dependent manner: the mean rate of occurrence was significantly reduced to 0.21±0.43 events·s^−1^·cell^−1^ and to 0.016±0.060 events·s^−1^·cell^−1^ by 30 nM and 100 nM Ryanozole, respectively (Fig. 4B, C). Time-based scan images revealed that, in the K4750Q cardiomyocytes showing isoproterenol-induced Ca^2+^ sparks (Fig. 4D, E) (Video S3), Ryanozole (100 nM) strongly suppressed the occurrence of Ca^2+^ sparks as well as Ca^2+^ waves and triggered activities (Fig. 4D, E) (Video S4).

### Ryanozole suppresses induced arrhythmias in K4750Q mice

We next tested the in vivo effect of Ryanozole on the K4750Q mice. In contrast to the R420W mice, ventricular arrhythmias could be induced in the K4750Q mice by adrenaline alone without caffeine. The experimental protocol was basically the same as that for R420W except that adrenaline alone was used instead of the Ad/Caf mixture. Typical ECG records are shown in Fig. 5A. Ventricular arrhythmias were often observed in the vehicle-pretreated mice (Fig. 5A, top). These arrhythmias were largely suppressed in the Ryanozole-pretreated mice (Fig. 5A, bottom). Ryanozole dose-dependently suppressed the rate of arrhythmias and T-wave alternans at 3 mg/kg or higher (Fig. 5B). At 30 mg/kg Ryanozole, almost no arrhythmias were induced by adrenaline administration.

**Fig. 5.**
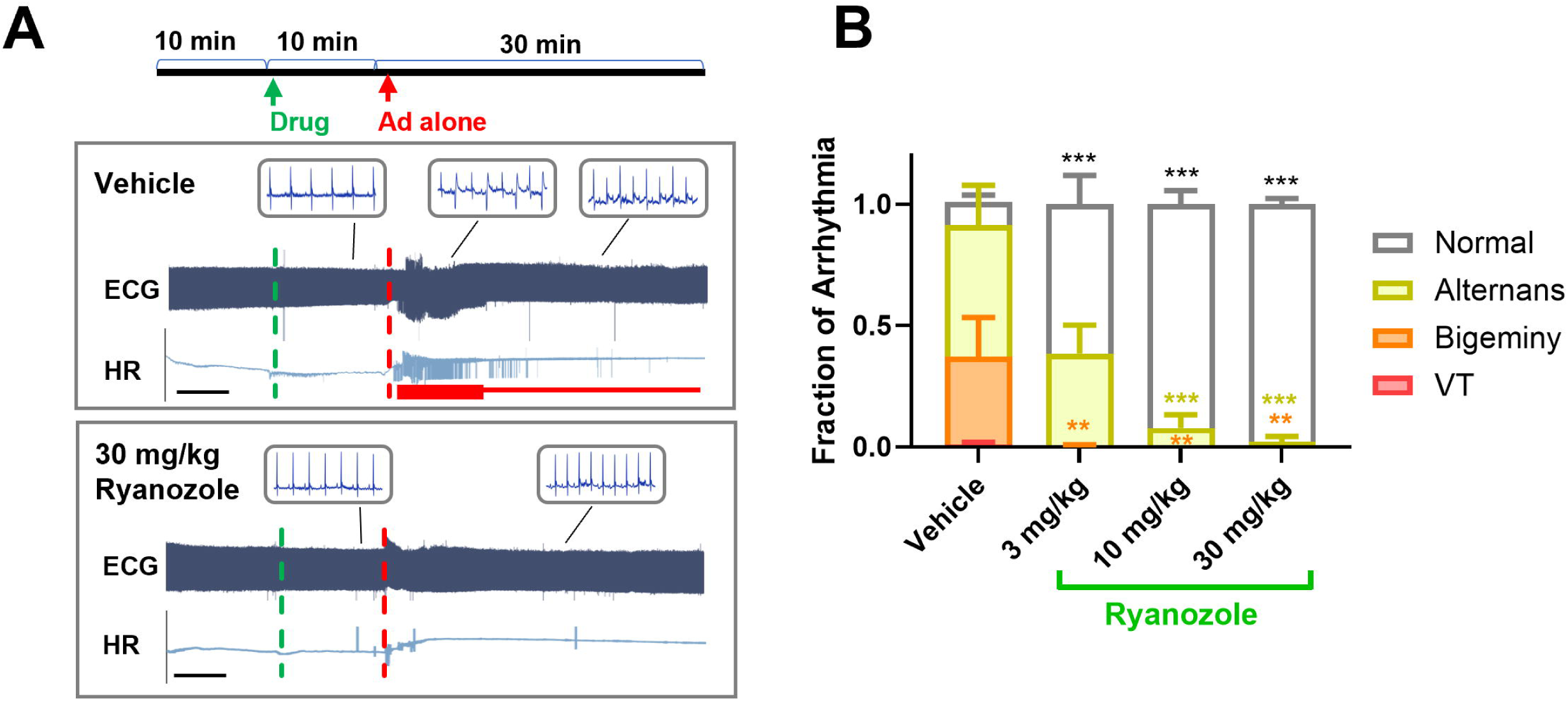
Effects of Ryanozole on induced arrhythmias in K4750Q mice. **A**, Typical effects of pretreated vehicle (top) and Ryanozole (bottom) on adrenaline (Ad)-induced arrhythmias. Ventricular arrhythmias are indicated with a horizontal thick red line, whereas T-wave alternans are indicated with a thin red line. Black bars indicate 5 min. **B**, Proportion of arrhythmias during 30 min after injection of Ad after pretreatment with Ryanozole (N=6−7). Data are represented as mean±SEM. **P<0.01, ***P<0.001 compared with Vehicle. Statistical significance was analyzed by two-way ANOVA followed by Dunnett’s multiple comparison test.

### Ryanozole suppresses activity-dependent arrhythmias in K4750Q mice

The *RYR2*-K4750Q variant has been reported to cause frequent polymorphic ventricular tachycardia in daily life (Kawamura et al., 2013). We therefore examined whether K4750Q mice also show spontaneous arrhythmias during daily activities. ECG transmitters were implanted subcutaneously in mice at 2–3 months of age and continuous ECG recordings were obtained up to 5 months of age. WT and R420W mice showed no arrhythmias, but all K4750Q mice showed intermittent ventricular arrhythmias, especially at night (Fig. 6A).

**Fig. 6.**
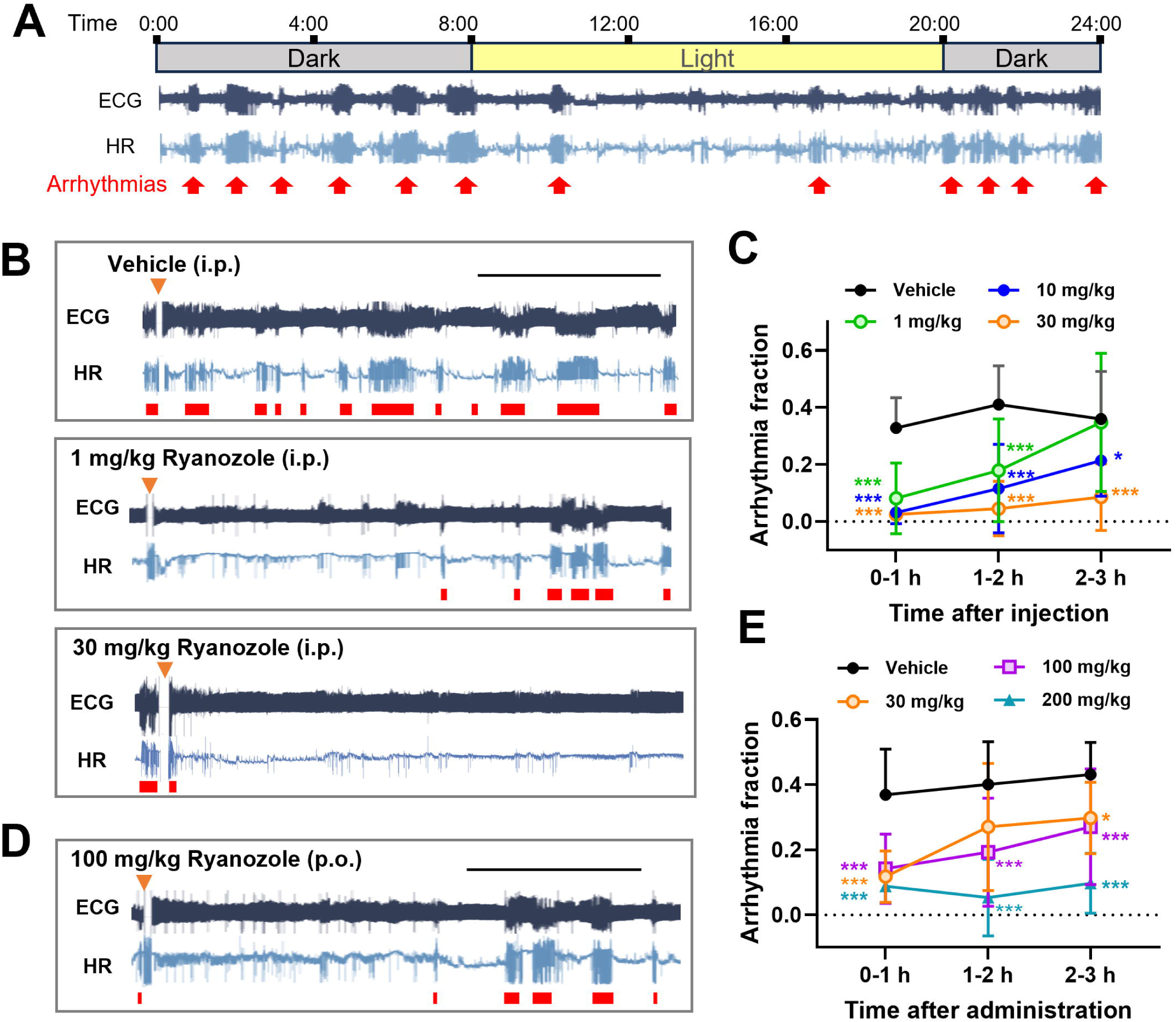
Effects of Ryanozole on spontaneous arrhythmias in K4750Q mice. **A**, Typical 24-hour ECG record. Red arrows indicate sustained ventricular arrhythmias. **B**, Typical ECG records after the intraperitoneal injection of vehicle (top), and Ryanozole at 1 mg/kg (middle) and 30 mg/kg (bottom). **C**, Average proportion of ventricular arrhythmia after the intraperitoneal injection of Ryanozole (n=8-10, N=4). **D**, Typical ECG records after the oral administration of vehicle (top) and Ryanozole (100 mg/kg). **E**, Average proportion of ventricular arrhythmia after the oral administration of Ryanozole (n=7-18, N=4). In **B** and **D**, thin black bar indicates 1 h, and thick red bars indicate ventricular arrhythmias. In E and G, data are represented as mean±SD. ;P<0.05, **P<0.01, ***P<0.001 compared with Vehicle. Statistical significance was analyzed by two-way ANOVA followed by Dunnett’s multiple comparison test.

We tested the effect of Ryanozole on spontaneous arrhythmias using RyR2-K4750Q mice. Because mice were active at night and showed more frequent arrhythmias, vehicle or Ryanozole was injected in the evening, between 7 and 8 p.m., and ECG records were analyzed for 3 h thereafter. Individual mice were administered various doses of Ryanozole or vehicle in a random order within 3 weeks. While numerous ventricular arrhythmias were observed after the injection of vehicle (Fig. 6B top), they were substantially suppressed after injection of 1 mg/kg (Fig. 6B middle) and almost completely disappeared with 30 mg/kg Ryanozole (Fig. 6B, bottom). On average, the rate of ventricular arrhythmia was significantly reduced with 1 mg/kg Ryanozole or more for at least 2 h (Fig. 6C). Although the suppressive effect of Ryanozole diminished over time, a significant effect still remained after 3 h at 30 mg/kg. Importantly, Ryanozole was also effective upon oral administration, for which higher doses were required to obtain effective suppression (Fig. 6D, E).

### Ryanozole does not impair cardiac function

Conventional antiarrhythmic drugs used for CPVT, i.e., β-blockers, Na^+^ channel blockers, and Ca^2+^ channel blockers, may be associated with side effects, including conduction disturbances, bradycardia, or reduced cardiac contractility, due to their own mechanism of action. To determine whether Ryanozole affects cardiac function, which suppresses RyR2 activity, we examined its effects on normal Ca^2+^ transients in cardiomyocytes and ECG parameters and echocardiographic parameters in vivo in WT and K4750Q mice.

Action potential-evoked Ca^2+^ transients were recorded in ventricular myocytes stimulated at 0.33 Hz in the presence of various concentrations of Ryanozole (Fig. 7A). In WT cells, peak F/F_0_ was significantly decreased, whereas 50% decay time was significantly increased by Ryanozole at 100 and 300 nM, but not at 30 nM (Fig. 7B, top). The effects of Ryanozole were fully reversed by washout of the drug. In RyR2-K4750Q cells, in contrast, no significant change in either peak F/F_0_ or 50% decay time was observed (Fig. 7B, bottom). In WT cells, it is likely that inhibition of RyR2 by Ryanozole at 100 nM or higher leads to a decrease in peak amplitude, which then causes prolonged decay of Ca^2+^ transients via the attenuation of Ca^2+^-dependent inactivation of the L-type Ca^2+^ channel. In RyR2-K4750Q cells, inhibition of leaky RyR2 could result in an increase in the SR Ca^2+^ content, which could mask the inhibitory action of the drug on RyR2.

**Fig. 7.**
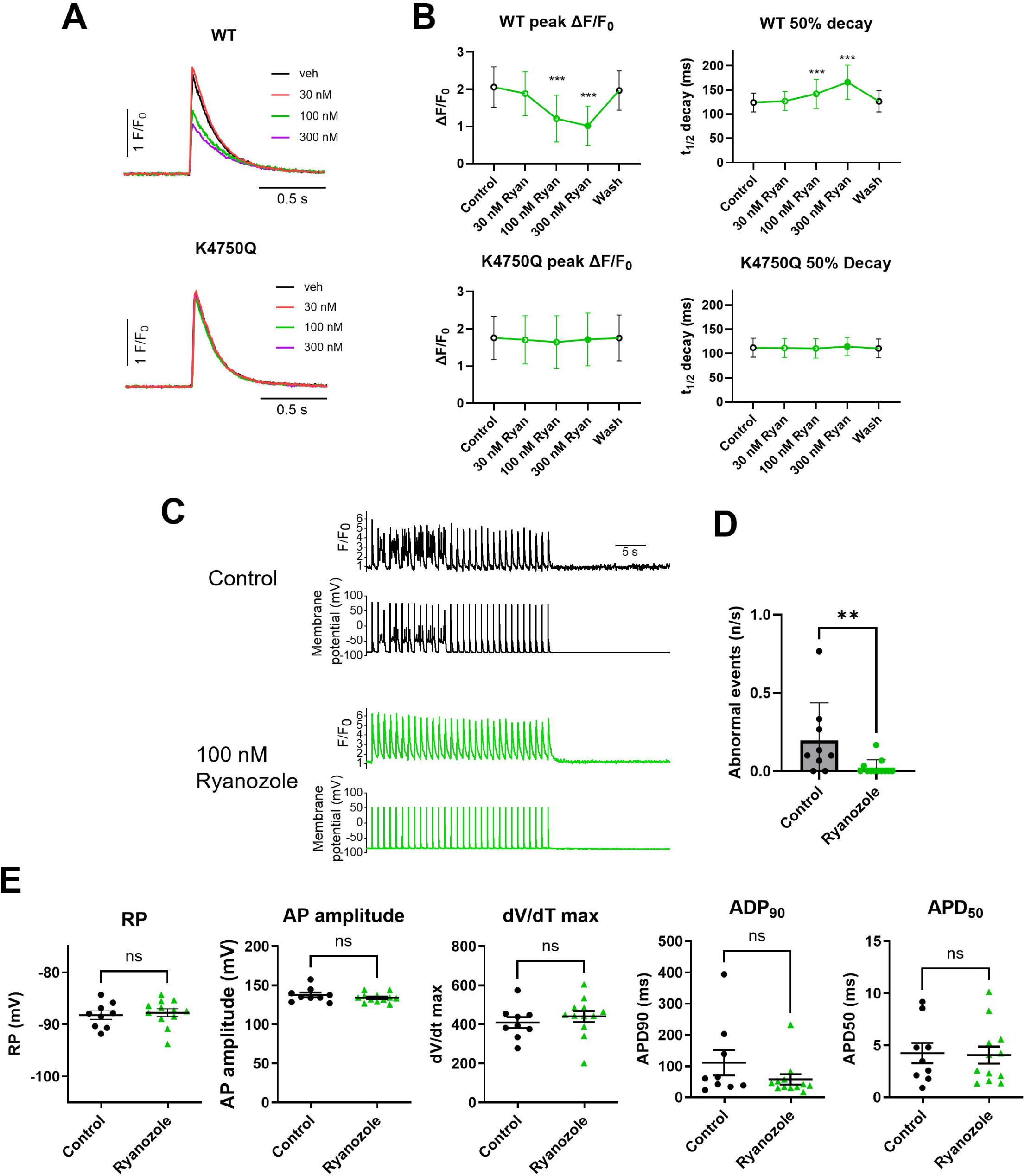
Effects of Ryanozole on Ca^2+^ transients and action potential characteristics in isolated cardiomyocytes from K4750Q mice. **A**, Typical Ca^2+^ transients (detected from ROIs) field-stimulated at 0.33 Hz in the absence and presence of Ryanozole in isolated cardiomyocytes from WT (upper) and K4750Q mice (lower). **B**, Effects of Ryanozole on average peak amplitude and 50 % decay time of Ca^2+^ transients in cardiomyocytes (n=84−86, N=4). **C**, Representative Ca^2+^ and membrane potential records in K4750Q cardiomyocytes in the absence and presence of Ryanozole. Action potentials were triggered by a 1 ms depolarizing current injection at 1 Hz 30 times under current clamp mode. **D**,. Effects of Ryanozole on frequencies of abnormal membrane potential signals (n=9–12). Data are mean±SD. Statistical significance was analyzed by Mann−Whitney test. ** p < 0.01 compared to Control. **E**, Effects of Ryanozole (100 nM) on resting potential (RP), action potential (AP) amplitude, dV/dt max, and action potential durations at 90% and 50% repolarization (APD_90_ and APD_50_, respectively) (n=9–12). Data are mean±SEM. Statistical significance was analyzed by unpaired t-test.

We also examined the effects of Ryanozole on action potential characteristics in cardiomyocytes from K4750Q mice by simultaneously monitoring membrane potentials and Ca^2+^ signals (Fig. 7C-E). Abnormal membrane potentials together with abnormal Ca^2+^ signals were frequently observed in K4750Q cardiomyocytes (Fig. 7C, top). In the presence of 100 nM Ryanozole, such abnormal events were significantly suppressed (Fig. 7D, bottom). Fig. 7E compares the action potential parameters. Resting potentials (RP), action potential (AP) amplitudes, dV/dt max, and action potential durations (APD) at 90% and 50% repolarization (APD_90_ and APD_50_, respectively) were rarely affected by Ryanozole at 100 nM. Notably, mean APD_90_ was shortened by half by the application of Ryanozole (from 111.0 ± 121.4 ms to 57.7 ± 57.4 ms, p = 0.195), although this did not reach significance. This can be explained by the suppression of Ca^2+^ release by Ryanozole, which results in reduction in inward NCX current at the late repolarization phase.

Figure 8A shows typical ECG traces before and after the intraperitoneal injection of vehicle or Ryanozole. Relative changes in ECG parameters, namely, heart rate, PR interval, and QRS interval, after drug administration are plotted in Fig. 8B. There are no significant differences in the heart rate, PR interval, and QRS interval between vehicle and 30 mg/kg Ryanozole administration. Fig. S4 shows the effects of typical conventional drugs, flecainide (20 mg/kg), L-type Ca^2+^ channel blocker verapamil (2.5 mg/kg), and β_1_-selective blocker atenolol (2 mg/kg) on ECG. The doses of these drugs were selected based on previous reports (Katz et al., 2010; Poll et al., 2021; Watanabe et al., 2009). These drugs actually significantly reduced the frequency of induced ventricular tachycardia in R420W mice (Fig. S4A) and of spontaneous arrhythmias in K4750Q mice (Fig. S4B). At effective doses, these drugs affected ECG parameters to varying degrees (Fig. S4C, D). Flecainide substantially prolonged PR and QRS intervals and reduced heart rate (Fig. S4C, D), verapamil slightly but significantly lengthened PR intervals (Fig. S4D, middle), and atenolol significantly reduced heart rate (Fig. S4D, left). These findings suggest that Ryanozole could provide antiarrhythmic effects with minimal alterations in ECG parameters, potentially offering an advantage over some existing therapies.

**Fig. 8.**
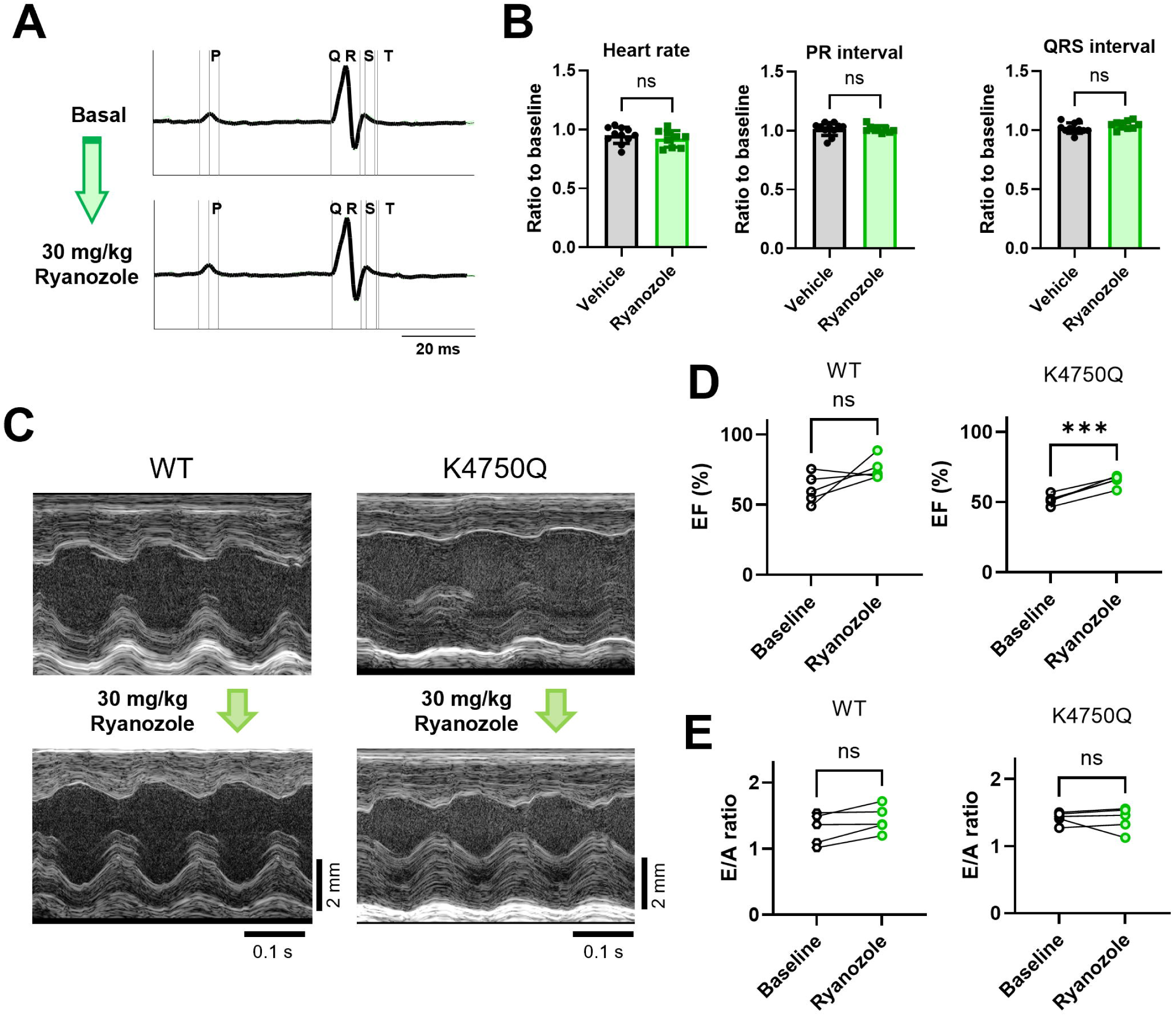
Effects of Ryanozole on cardiac functional properties. **A**, Typical ECG wave forms before and after the intraperitoneal injection of Ryanozole (30 mg/kg). **B**, Average ECG parameters before and after the intraperitoneal injection of Ryanozole (N=10, 5 R420W and 5 K4750Q mice). Statistical significance was analyzed by Mann−Whitney test. **C**, Representative images of M-mode echocardiography from WT mice (left) and K4750Q (right) before and after the intraperitoneal injection of Ryanozole. **D and E**, Graphs showing the data from echocardiography regarding left ventricular ejection fraction (EF) (**D**) and E/A ratios (**E**) before and after the intraperitoneal injection of Ryanozole (N=5). Statistical significance was analyzed by paired t-test.

We also assessed the effects of Ryanozole on cardiac function by echocardiography (Fig. 8C, D) in WT and K4750Q mice. Intraperitoneal injection of Ryanozole did not change the ejection fraction (EF) in WT mice (Fig. 8C, left and 8D, left). Interestingly, Ryanozole significantly enhanced EF in K4750Q mice (Fig. 8C, right and 8D right), in which the basal EF was marginally lower than in the WT (Fig. 8D). In addition, Ryanozole preserved the diastolic index “early (E)/late (A) mitral inflow peak velocity ratio” (E/A) in WT and K4750Q mice (Fig. 8E). Taken together, these findings suggest that Ryanozole may provide antiarrhythmic benefits with a favorable cardiac safety profile, including minimal alterations in cardiac function or ECG parameters.

## Discussion

In this study, we explored the therapeutic potential of the RyR2-specific stabilizer Ryanozole on ventricular arrhythmias using two lines of CPVT-associated mutant RyR2 mice. Our findings indicate that Ryanozole preferentially suppresses Ca^2+^ release at low [Ca^2+^]_cyt_ and effectively inhibits arrhythmogenic spontaneous Ca^2+^ sparks and Ca^2+^ waves without significantly affecting physiological Ca^2+^ transients. We demonstrated that Ryanozole is very effective in preventing catecholamine-induced arrhythmias and stopping spontaneous arrhythmias during daily activity. Importantly, Ryanozole did not impair cardiac function at its effective doses, suggesting its potential as a safe and effective therapeutic candidate.

It has been postulated that drugs with RyR2-stabilizing action have an antiarrhythmic effect on CPVT and other ectopic arrhythmias (Do & Knollmann, 2025; Svensson et al., 2023). Actually, several compounds/drugs with RyR2-modulating effects have been reported to have antiarrhythmic activity to support the idea that RyR2 is a promising target for the treatment of arrhythmias. EL20, an analog of tetracaine (Klipp et al., 2018; Word et al., 2021), and the unnatural verticilide (Batiste et al., 2019) have been reported to suppress RyR2 and arrhythmias in arrhythmic model mice. A group of compounds such as Rycal drugs that stabilize the association of FK binding protein 12.6 (FKBP12.6/calstabin2) with RyR2 has also been reported to ameliorate abnormal Ca^2+^ release in cardiomyocytes (Andersson & Marks, 2010; Marks, 2023). In addition, flecainide is an effective antiarrhythmic drug that mainly targets Na^+^ channels, and its action on RyR2 has been reported to contribute to its antiarrhythmic effect (Bannister, MacLeod, & George, 2022; Salvage, Huang, Fraser, & Dulhunty, 2022; Watanabe et al., 2009). Meanwhile, a β-blocker, carvedilol, has also been reported to suppress abnormal Ca^2+^ release via RyR2 (Xiao et al., 2016; Zhou et al., 2011).,

Ryanozole is a compound with high affinity and specificity for RyR2 (Ishida et al., 2023). According to the unaltered ECG records, it does not appear to act on major ion (Na^+^, K^+^, or Ca^2+^) channels in the mouse heart. The modulating effects of Ryanozole are not mediated by the associated protein, FKBP12.6, because they inhibited FKBP12.6-unbound RyR2 in HEK293 cells that do not express FKBP12.6. Ryanozole also appears to suppress both calmodulin-unbound and calmodulin-bound RyR2, as the amount of calmodulin in RyR2-expressing HEK293 cells is insufficient in terms of the molar ratio compared with the amount of RyR2 (Gao et al., 2023). It has been reported that RyR2 is subject to post-translational modifications such as phosphorylation (Huke & Bers, 2008; Lanner et al., 2010; Potenza et al., 2019; Takenaka et al., 2023). However, the effect of Ryanozole was indistinguishable between the phospho-null triple mutant (S3A) and phospho-mimetic triple mutant (S3D), suggesting its phosphorylation-independent action (Fig. S1G, H). Thus, Ryanozole may directly target RyR2 to stabilize its activity in the closed state. Importantly, Ryanozole similarly modulates RyR2 carrying four different CPVT mutations, which are localized in the N-terminal, central, and C-terminal regions (Fig. S1C–F). These findings suggest that Ryanozole may be effective against a variety of RyR2 mutations found in CPVT.

Ryanozole strongly suppressed the occurrence of Ca^2+^ waves and Ca^2+^ sparks with relatively minor effects, if any, on action potential-evoked Ca^2+^ transients in isolated cardiomyocytes from CPVT model mice (Fig. 2B, Fig. 4C). This may be related to the virtual absence of effects of Ryanozole on cardiac function (Fig. 6E, F). This unique property may be explained by the mechanism of action of the compound: Ryanozole strongly suppresses RyR2 activity at low [Ca^2+^]_cyt_, while mildly suppressing maximal activity at high [Ca^2+^]_cyt_ (Fig. 1D). As a result, the generation of local Ca^2+^ waves and Ca^2+^ sparks at resting [Ca^2+^]_cyt_ may be effectively suppressed, but action potential-evoked Ca^2+^ transients caused by massive Ca^2+^ influx via L-type Ca^2+^ channels may be less suppressed. Similar results were also obtained with other RyR2 inhibitors such as chloroxylenol and riluzole (Takenaka et al., 2023). They also reduced the affinity for activating Ca^2+^ of [^3^H]ryanodine binding and mildly suppressed peak activity, and also effectively suppressed the occurrence of Ca^2+^ waves and Ca^2+^ sparks with minor effects on action potential-evoked Ca^2+^ transients. Identification of the binding site and the resulting conformational changes in RyR2 would provide critical insights into the underlying mechanism of RyR2 specificity and Ca^2+^-dependent inhibitory action of Ryanozole.

In addition to the findings on Ryanozole, this study provides insights into the procedure for testing drug effects using CPVT model mice. We employed two RyR2 mutant lines with different arrhythmia severity. Mice with homozygous RyR2-R420W mutation show a moderate phenotype and require strong stimulation (i.e., catecholamine plus caffeine) to exhibit arrhythmias, as in typical CPVT model mice (Cerrone et al., 2005; Kryshtal et al., 2021; Loaiza et al., 2013; Radwanski et al., 2015). A mutation at the same site, R420Q, is also known to cause familial CPVT (Domingo et al., 2015). We also generated RyR2-K4750Q mice, which have a more severe phenotype than R420W, exhibiting spontaneous arrhythmias during daily activity. We previously reported that RyR2-I4093V mice, which were created by a random mutation, showed numerous spontaneous arrhythmias and are useful for assessing antiarrhythmic effects (Okabe et al., 2024). RyR2-K4750Q mice have milder arrhythmia symptoms than I4093V mice, but still exhibit spontaneous intermittent arrhythmias. Thus, mice harboring highly augmented RyR2 mutations may have the propensity to suffer spontaneous arrhythmias and appear to be very useful for evaluating the arrhythmia-blocking effect of drugs and the time course of drug effects.

In addition to preventing catecholamine-induced arrhythmias, Ryanozole also reduced ongoing arrhythmias (Fig 6), suggesting a potential for acute therapeutic use. The fact that there were no changes in ECG parameters or cardiac function suggests that Ryanozole is a promising candidate as a new class of antiarrhythmic drug. Because Ryanozole did not suppress cardiac function in the WT and rather improved it in K4750Q mice, it may have advantages over conventional drugs in patients with heart failure. In addition, this important property raises the possibility that Ryanozole may also be effective against arrhythmias in CPVT caused by mutations in proteins other than RyR2 such as calsequestrin 2 (CASQ2) (Faggioni & Knollmann, 2012; Priori & Chen, 2011) and calmodulin (Gao et al., 2023), and heart failure itself. In particular, increased RyR2 leakage has been reported in CASQ2-linked CPVT, heart failure and ischemia-reperfusion (Andersson & Marks, 2010; Bers, Eisner, & Valdivia, 2003; Bovo, Mazurek, & Zima, 2018; Boyden & Smith, 2018; Eisner, Caldwell, Kistamas, & Trafford, 2017; Marks, 2023). For patients with impaired cardiac function, antiarrhythmic drugs that do not impair cardiac function would be safe and easy to use. However, further studies will be required to evaluate its efficacy in these settings.

In conclusion, our findings highlight Ryanozole as a promising RyR2-specific modulator with potent antiarrhythmic activity and minimal impact on cardiac function in CPVT mouse models. Future work will be needed to optimize its pharmacokinetic properties and assess its efficacy and safety in large animal models and clinical settings.

## Author Contributions

N.K., M.Kod, T.M, M.Kon, M.S., and H.I., designed the study. N.K., M.Kod, T.M, M.Kon, M.S., H.I., K.I., and Y.E. performed functional analyses. R.I., S.M., X.Z., and H.K. synthesized and provided Ryanozole. Y.U.I., T.I., S.N., and H.N. generated and provided mouse models. N.K., M.Kod, T.M., M.Kon, M.S., H.I., R.I., K.I., S.M., Y.E., U.Y., J.K., H.K., and T.S. interpreted the data. N.K., T.M., J.K., and H.K. conceived the project and acquired the critical funding for this project (DNW-21012, AMED). N.K., M.Kod, T.M., and M.Kon wrote the manuscript. All authors reviewed, edited, and approved the manuscript.

## Acknowledgments

We thank Mirei Takahashi and Ikue Hiraga for technical assistance. We also thank staff at the Center for Biomedical Research Resources and the Laboratory of Proteomics and Biomolecular Science, Research Support Center, Juntendo University Graduate School of Medicine, for technical assistance. We are also grateful to Edanz (https://jp.edanz.com/ac) for editing a draft of this manuscript.

## DECLARATION OF TRANSPARENCY AND SCIENTIFIC RIGOUR

