## Supplemental Figures 1-4 for "A novel RyR2-selective stabilizer prevents stress-induced ventricular arrhythmias without impairing cardiac function"

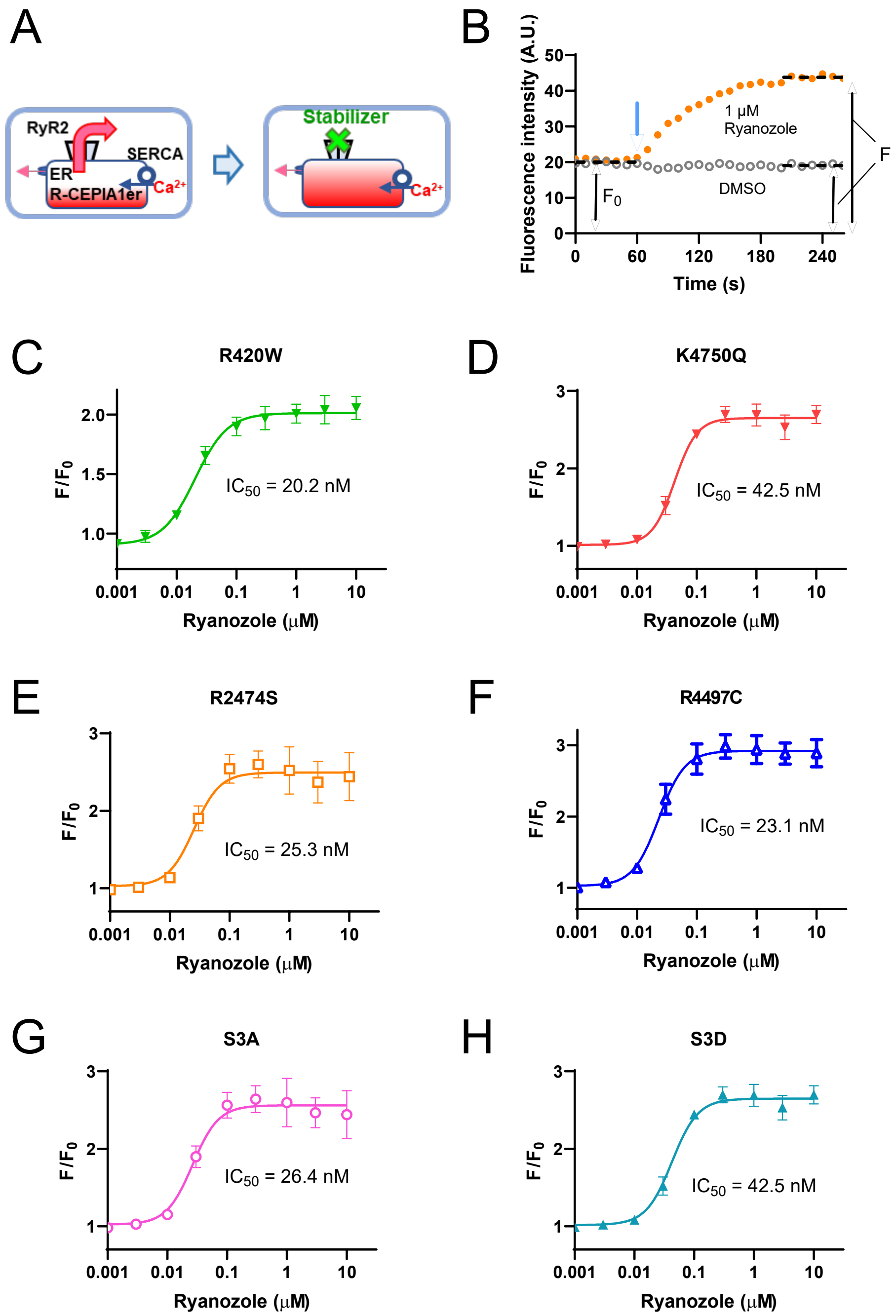


**Fig. S1: Effects of Ryanozole on F/$F_0$ signals in HEK293 cells expressing R-CEPIA1er and mutant RyR2s.**

**A,** Schematic of ER $Ca^{2+}$-based evaluation of RyR2 modulators using HEK293 cells expressing RyR2 and ER $Ca^{2+}$ sensor R-CEPIA1er. **B,** Representative time-lapse R-CEPIA1er fluorescence measurement using a FlexStation3 fluorometer. Test solutions containing DMSO (0.2%) and Ryanozole (1 μM) were added at 60 s (blue arrow) and ratios (F/$F_0$) of fluorescence intensity before ($F_0$) and after addition of the compounds (F) were obtained as an indicator of RyR2 inhibition. **C–H,** Dose-dependent effects of Ryanozole on F/$F_0$ signals in HEK293 cells expressing R420W (**C**), K4750Q (**D**), R2474S (**E**), R4497C (**F**), phospho-null triple mutant of RyR2, S2807A/S2813A/S2030A (RyR2 S3A) (**G**), and phospho-mimetic triple mutant of RyR2, S2807D/S2813D/S2030D (RyR2 S3D) (**H**).

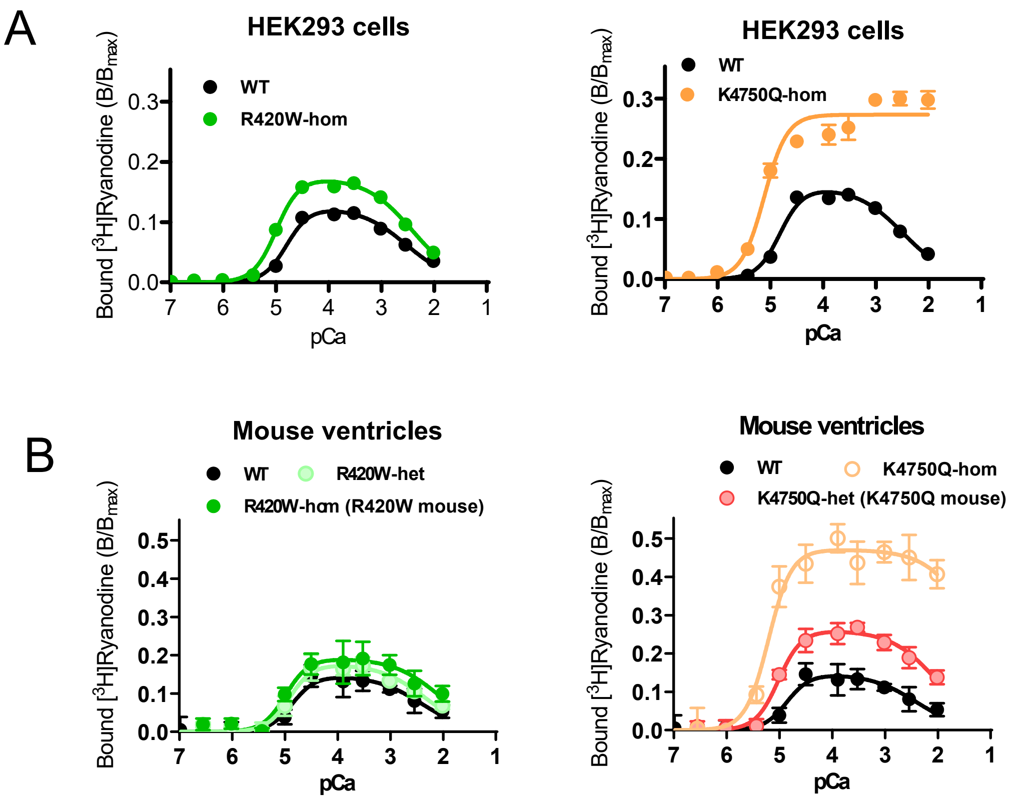


**Fig. S2: Channel activities of R420W and K4750Q determined using [$^3$H]ryanodine binding.**

**A**, [$^3$H]Ryanodine binding of homozygous RyR2-R420W (left) and homozygous K4750Q (right) expressed in HEK293 cells. **B**, [$^3$H]Ryanodine binding of ventricular microsomes from R420W (left) and K4750Q mice (right). R420W-het: heterozygous R420W; R420W-hom: homozygous R420W; K4750Q-het: heterozygous K4750Q; K4750Q-hom: homozygous K4750Q. Note that the data for R420W-hom and K4750Q-het are the same as those in Fig. 4A.

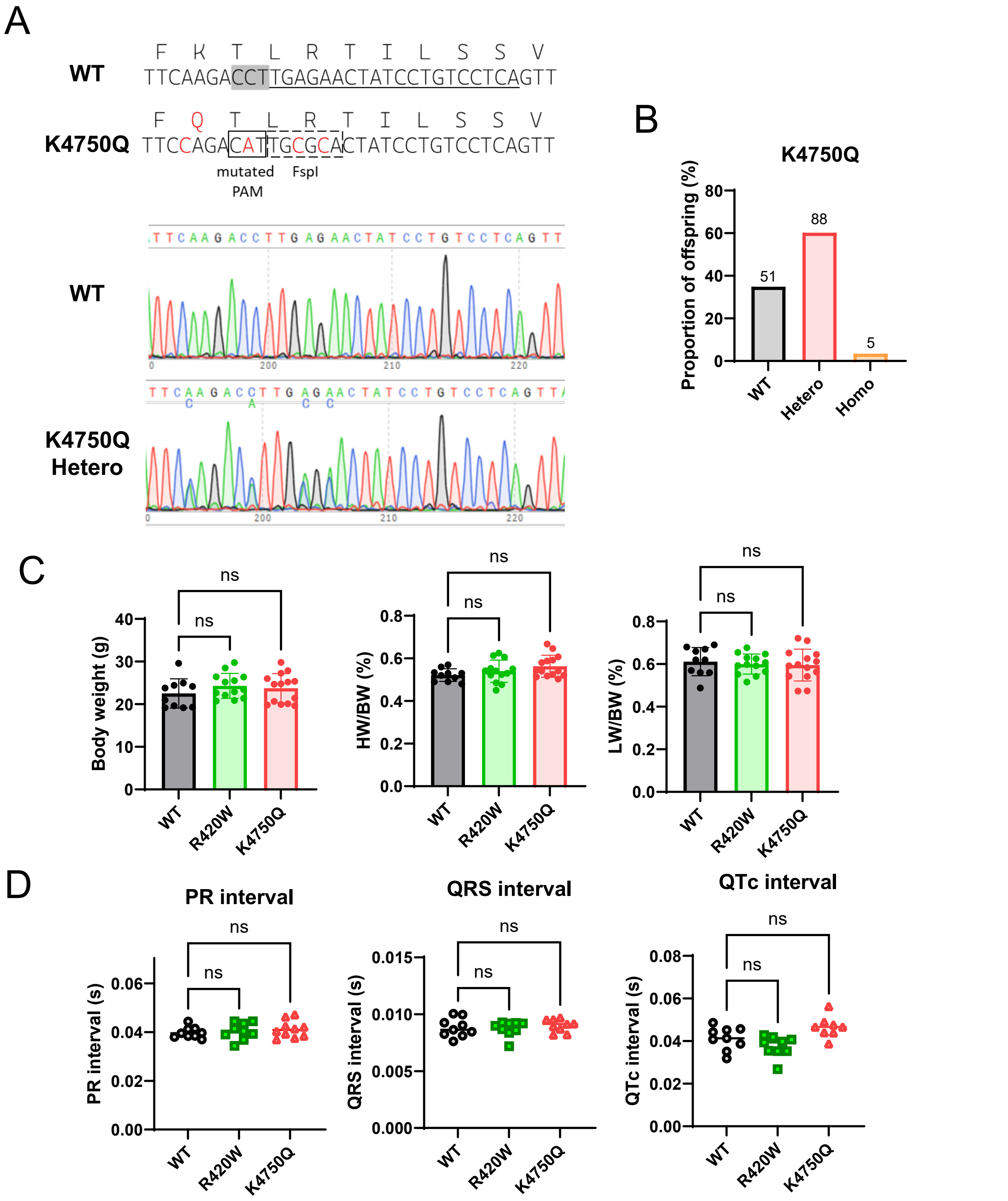


**Fig. S3: Generation of knock-in mice harboring RyR2-K4750Q mutation and comparison of in vivo data among WT, R420W, and K4750Q mice.**

**A,** The sequences of WT and K4750Q mutant alleles (top) and the sequences of an amplified DNA fragment from tails of WT and heterozygous (Hetero) offspring (bottom). K4750Q was generated by CRISPR/Cas9 using single-stranded donor oligonucleotide carrying Fsp I restriction site (TGCGCA). The sequence of exon 99 of the RYR2 gene (underlined) was selected for crRNA synthesis. The PAM next to the sequence of guide RNA is indicated by shade. **B,** Genotype ratio of offspring produced by heterozygous K4750Q mating. **C,** Body weight (left), heart to body weight ratio (middle), and lung to body weight ratio (right) of WT, R420W, and K4750Q mice at 2–3 months old. **D,** ECG parameters of WT, R420W, and K4750Q mice at 2–3 months old. R420W and K4750Q in **C** and **D** refer to homozygous R420W and heterozygous K4750Q mice, respectively. In **C**-**D**, statistical significance was analyzed by one-way ANOVA followed by Dunnett's multiple comparison test.

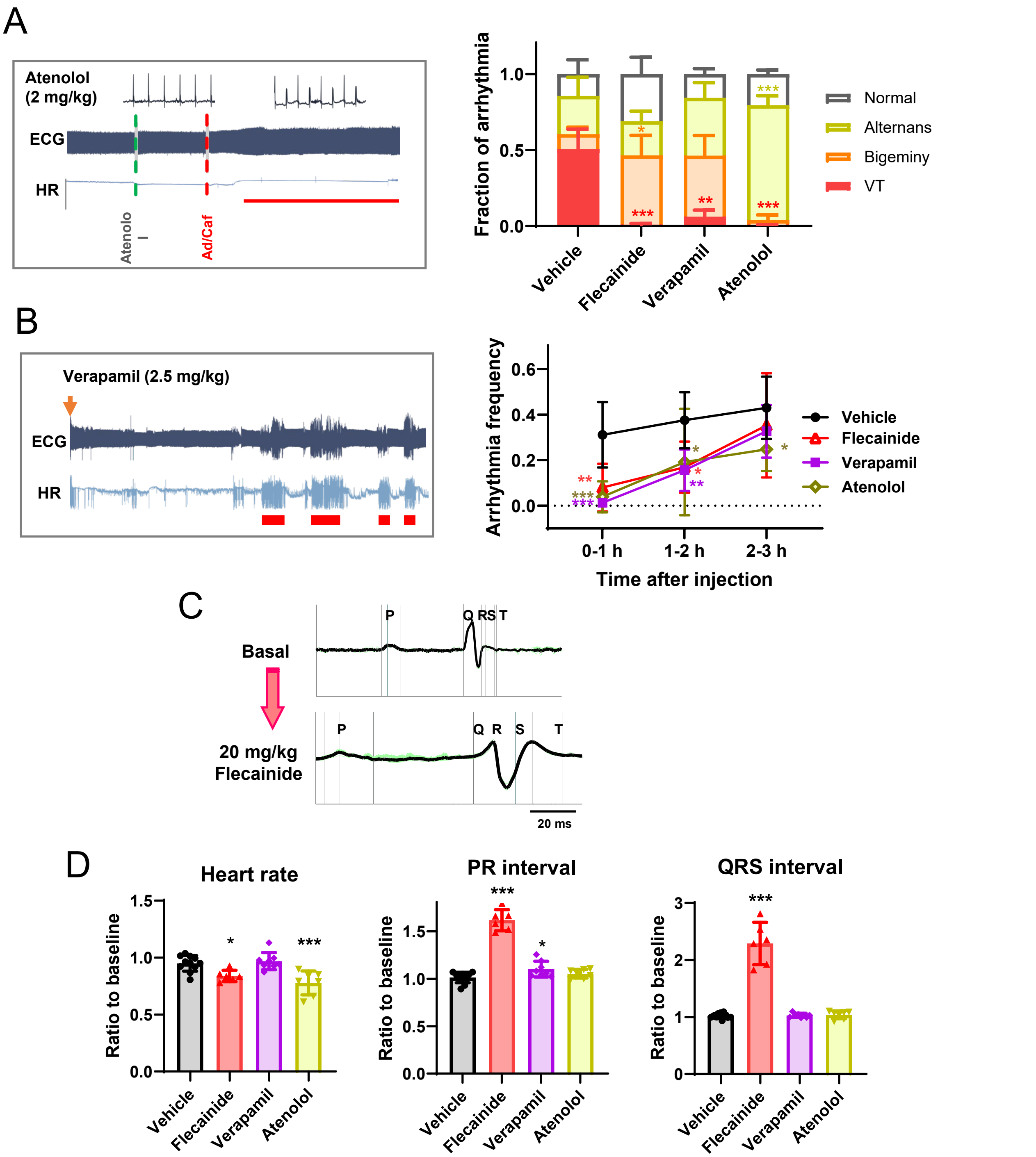


**Fig. S4 : Effects of conventional antiarrhythmic drugs on induced and spontaneous arrhythmias and ECG parameters in mutant RyR2 mice.**

**A,** Effects of drugs on Ad/Caf-induced arrhythmias in R420W mice. Left: typical effect of atenolol. A thin red line indicates T-wave alternans. Right: averaged effects of flecainide (20 mg/kg), verapamil (2.5 mg/kg), and atenolol (2 mg/kg) (n=6–8, N=4). Data are represented as mean ± SEM. **P<0.01, ***P<0.001 compared with Vehicle. Statistical significance was analyzed by two-way ANOVA followed by Dunnett's multiple comparison test. **B,** Effects of drugs on spontaneous arrhythmias in K4750Q mice. Left: typical effect of verapamil. Thick red bars indicate ventricular arrhythmias. Right: averaged effects of drugs (right) are shown (n=6–8, N=4). Doses of drugs were the same as those in **A**. Data are represented as mean ± SD. **P<0.01, ***P<0.001 compared with Vehicle. Statistical significance was analyzed by two-way ANOVA followed by Dunnett's multiple comparison test. **C,** Representative ECG waveforms before and 10 min after the intraperitoneal administration of flecainide. **D,** Comparison of ECG parameters before and after injection of drugs (N=10, 5 R420W and 5 K4750Q mice). Doses of drugs were the same as those in **A**. Data are mean ± SD. *P< 0.05, ***P<0.001 compared with Vehicle. Statistical significance was analyzed by one-way ANOVA followed by Dunnett's multiple comparison test.
